# Measuring Connectivity in Linear Multivariate Processes with Penalized Regression Techniques

**DOI:** 10.1101/2023.07.17.549308

**Authors:** Y. Antonacci, J. Toppi, A. Pietrabissa, A. Anzolin, L. Astolfi

**Affiliations:** Dipartimento di Ingegneria, Universitá di Palermo, Italy; Department of Computer, Control and Management Engineering, University of Rome La Sapienza, Rome, Italy; IRCCS Fondazione Santa Lucia, Rome, Italy; Spaulding Rehabilitation Hospital, Harvard Medical School, Boston, MA, USA; Athinoula A. Martinos Center for Biomedical Imaging, Massachusetts General Hospital, Harvard Medical School, Boston, MA, USA

## Abstract

The evaluation of time and frequency domain measures of coupling and causality relies on the parametric representation of linear multivariate processes. The study of temporal dependencies among time series is based on the identification of a Vector Autoregressive model. This procedure is pursued through the definition of a regression problem solved by means of Ordinary Least Squares (OLS) estimator. However, its accuracy is strongly influenced by the lack of data points and a stable solution is not always guaranteed. To overcome this issue, it is possible to use penalized regression techniques. The aim of this work is to compare the behavior of OLS with different penalized regression methods used for a connectivity analysis in different experimental conditions. Bias, accuracy in the reconstruction of network structure and computational time were used for this purpose. Different penalized regressions were tested by means of simulated data implementing different ground-truth networks under different amounts of data samples available. Then, the approaches were applied to real electroencephalographic signals (EEG) recorded from a healthy volunteer performing a motor imagery task. Penalized regressions outperform OLS in simulation settings when few data samples are available. The application on real EEG data showed how it is possible to use features extracted from brain networks for discriminating between two tasks even in conditions of data paucity. Penalized regression techniques can be used for brain connectivity estimation and can be exploited for the computation of all the connectivity estimators based on linearity assumption overcoming the limitations imposed by the classical OLS.

## INTRODUCTION

In the framework of linear signal processing, Vector Autoregressive model (VAR) has been proved to be a robust and reliable tool for analyzing temporal dependencies among time series in several research fields, ranging from economics to biomedical sciences. In neuroscience, these models are extensively used for evaluating brain connectivity in order to understand how different brain areas communicate [1]. With the term brain connectivity, we refer both to the concept of “functional connectivity”, that is the evaluation of statistical dependencies between different neuronal units, and “effective connectivity” that describes networks with directional effects, thereby requiring the generation of a mechanistic model of the cause-effect relationships [2]. Granger causality (GC) is a versatile tool for the analysis of the cause-effect relationships between different time series and was firstly formulated in the framework of bivariate autoregressive modeling, stating that a time series *X* G-causes another time series *Y*if the past of *X* contains information that helps to predict the future of *Y* above and beyond the information already contained in the past of *Y*[3]. To account for the influence of other time series, the bi-variate formulation has been extended to the multivariate case by using multivariate autoregressive (MVAR) models, leading to the well-known conditional form of the GC [4]. The MVAR analysis allows the time and frequency domain representation of physiological time series to be derived directly from model coefficients and their frequency-domain representation [5]. Accordingly, several measures of connectivity have been introduced and applied in recent years, many of which are based on the concept of GC and rely on the identification of the MVAR model. The most used measures, able to quantify causality in the frequency domain, are the directed transfer function (DTF) [6], the directed coherence (DC) [7] and the partial directed coherence (PDC) with all its formulations [8, 9]. These measures are widely used for the analysis of the interactions between physiological time series and, more generally to characterize brain connectivity [10].

The computation of all these connectivity estimators is based on the identification procedure of an MVAR model which in turn includes the solution of a linear regression problem to estimate the matrix of autoregressive (AR) coefficients and the innovation variance. The Ordinary Least Squares (OLS) method finds the optimal solution, starting from the hypothesis of independence between regressors and residuals of the regression problem [1]. However, its accuracy is strongly influenced by the number of available data samples. In particular, the mean squared error (MSE) in the MVAR parameters estimation decreases when the amount of available data increases [11]. As a rule of thumb, it is necessary that the ratio between the number of data samples available and the number of parameters to be estimated (k-ratio) should be at least equal to 10 to ensure the accuracy of the OLS estimator [12, 13]. Otherwise, particularly when the k-ratio is close to one, the estimation problem becomes ill-posed and underdetermined [12, 13]. In such a situation, the OLS does not guarantee the uniqueness of the solution, leading to an ambiguity in the MVAR representation of the data. One possible solution to overcome this methodological limitation, is to use penalized regression techniques. The idea is to add a constraint (based on *l*_1_-norm and/or *l*_2_-norm) to regularize the OLS problem. The effect of *l*_2_-norm is to shrink the MVAR parameters towards zero, while the effect of *l*_1_-norm is to select only specific coefficients, setting the others to zero. Both *l* _2_- and *l* _1_-norm methods can improve the OLS accuracy when the k-ratio is lower than 10, with the effect of also reducing the mean square error [13, 14]. Penalized regressions can be convex or non-convex depending on whether the constrained optimization problem they solve guarantees or not the existence of a global minimum [15]. In the literature they have been used for GC estimation [13, 16–19], neuro-imaging data analysis [20] and brain connectivity analysis between EEG time series in a single trial or in real-time settings [21, 22]. Nevertheless, these applications show some limitations: the comparison between different penalized regressions has been done only for a single value of k-ratio; for a real-time application, only one method based on *l*_1_-norm was used; convex and non-convex methods were directly compared even if it is well known that the latter do not achieve a stable solution. Although penalized regression techniques have been proposed in different studies, they are not commonly used to study brain connectivity from EEG signals. Therefore, this work is motivated by the fact that there are no available extensive studies that analyze the behavior of penalized regressions in both simulated and real EEG time series by also providing guidelines for their exploitation. Specifically, we have selected five penalized regression techniques that solve a linear convex optimization problem guaranteeing the existence of a solution: Ridge Regression (RR) [23], Least Absolute Shrinkage and Selecting Operator regression (LASSO) [24], Elastic-Net regression (E-NET) [25], fused LASSO regression (F-LASSO) [26] and Sparse group LASSO regression (SG-LASSO) [27]. We will aim to demonstrate by using both simulated and real EEG data, that it is possible, also in conditions of data paucity, to accurately reconstruct connectivity patterns describing networks of several coupled Gaussian systems exhibiting complex interactions, and to discriminate between two different experimental conditions by using features extracted from a brain connectivity analysis starting from short windows of EEG signals [21].

The remainder of this paper is organized as follows. The penalized regression techniques are presented in Sec.. Sec. presents a simulation study on EEG-like surrogate data, generated by reproducing ground-truth networks, for different values of the k-ratio. Sec., presents an application of penalized regression techniques to real EEG data, recorded from a healthy subject performing an motor imagery (MI) task. Finally, in Sec. the main features of the different regression techniques are discussed and conclusions are drawn.

## METHODS

In this section, we first introduce the MVAR model identification procedure. Then, we describe the different penalized regressions by highlighting their distinctive features and the underlying principles of each presented technique.

### MVAR model Identification

Let us consider a dynamical system 𝓎, whose activity is mapped by a discrete-time stationary vector stochastic process composed of *M* real-valued zero-mean stationary scalar processes, *Y* = [*Y*_1_ …*Y*_*M*_]. Considering the time step *n* as the current time, the present and the past of the process are denoted as *Y*_*n*_ = [*Y*_1,*n*_ …*Y*_*M*_,_*n*_] and 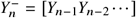, respectively. Moreover, assuming that *Y* is a Markov process of order *p*, its whole past history can be truncated using *p* time steps, i.e. using the *M p*-dimensional vector 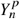 so that 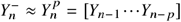. Then, in the linear signal processing framework, the dynamics of *Y* can be completely described by the VAR model:

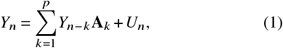

where **A**_*k*_ ∈ *ℜ*^*M*×*M*^ is the matrix containing the autoregressive (AR) coefficients, and *U* = [*U*_1_ …*U*_*M*_] ∈ *ℜ*^*M*×1^ is a zero-mean white process, denoted as innovations, with covariance matrix 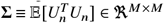 (𝔼 is the expectation value).

Let us now consider a realization of the process *Y* involving *N*_*s*_ consecutive time steps, collected in the data matrix 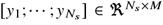, where the operator “;” stands for row separation, so that the *i* ^*th*^ row is a realization of *Y*_*i*_, i.e. *y*_*i*_ = [*y*_1,*i*_…*y*_*M,i*_], *i* = 1, …, *N*_*s*_, and the *j*^*th*^ column is the time series collecting all realizations of *Y*_*j*_, i.e. [*y* _*j*,1_…*y* _*j,Ns*_]^*T*^, *j* = 1, …, *M*. OLS finds an optimal solution for the problem (1) by solving the following linear quadratic problem [1]:

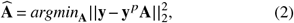

where 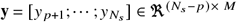 is the matrix of the responses values, 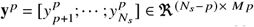 is the matrix of the regressors and **A** = [**A**_1_; …; **A**_*p*_] ∈ *ℜ*^*M p*×*M*^ is the matrix of coefficients. The problem has a solution in a closed form 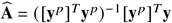 for which the residual sum of squares is minimized. When *N*_*s*_ − *p ≤ M p* the OLS does not guarantee the uniqueness of the solution since the matrix ([***y***^*p*^]^*T*^***y***^*p*^) becomes singular [15].

### Penalized linear regression techniques

From a mathematical point of view, regularizing the OLS problem means adding a constraint to the problem (2). In the Lagrangian form, the constraints are written in the cost function as additional weighted costs. In this scenario, it is possible to carry out separate regression analyses for each process composing the dynamical system Y. In other words, it is possible to estimate separately each column *A* _*j*_ of **A**. The multi-objective optimization problems reads as:

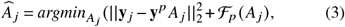

where 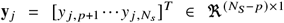 denotes the j-th column of **y** containing the response vector for the specific response variable (*j* = 1, …, *M*), *A* _*j*_ = [*a* _*j*,1_(1), …, *a* _*j,M*_ (1), …, *a* _*j*,1_(*p*), …, *a* _*j,M*_ (*p*)]^*T*^ ∈ *ℜ*^*M p*×1^ is the vector coefficient corresponding to the response variable *j* and ℱ_*p*_ is a penalty function applied to *A* _*j*_. As anticipated in Section, in this work we analyze the linear penalty functions reported in Table I.

**TABLE I.**
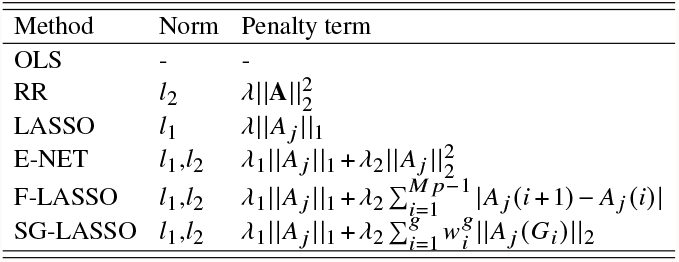
Penalty terms associated with problem 3.

RR is characterized by a *l*_2_-norm based term that shrinks the estimated AR parameters towards zero, reducing the MSE

[23]. The regularization parameter *λ* controls the amount of penalization to be applied, if *λ* = 0, RR reduces to OLS. The minimization of the RR functional leads to the closed form solution **Â**= [(***y***^*p*^)^*T*^***y***^*p*^ + **λI**]^−1^ (***y***^*p*^)^*T*^***y***, where **I**∈ℜ^*M*^ _*p*_ × ^*M*^ _*p*_ is the identity matrix. Besides the reduction of the MSE, it would be of interest to reduce the number of selected parameters (i.e., parameters where non-null values are imposed) especially when *N*_*s*_ *< M p*. This result could be achieved by using a penalty term based on the *l*_0_-norm, which in turn is equal to the number of non-zero elements in a vector [14]. Unfortunately, such a penalization function would render the optimization problem non-convex [28]. The effect of the *l*_0_-norm can be approximated by using penalty terms based on the *l*_1_-norm, such as the LASSO: the penalty term acts so as to shrink some coefficients and set others to 0 [24]. In the LASSO, the parameter *λ* sets the trade-off between the number of non-null coefficients selected in the matrix **A** and residuals sum of squares (RSS).

Similarly, the E-NET simultaneously performs the automatic variable selection and the coefficient shrinkage [25]. The variable selection is regulated by the parameter *λ*_1_, weighting the term based on the *l*_1_-norm. The shrinkage of coefficients, instead, is regulated by the parameter *λ*_2_ weighting the term based on the *l*_2_-norm. The E-NET coincides with RR if *λ*_1_ = 0 and with LASSO if *λ*_2_ = 0. F-LASSO includes in its penalty function a further term with respect to LASSO, which computes the *l*_1_-norm of the vector of the differences between the coefficients of successive predictors. This term is used to enforce smoothness along the predictors, i.e. along the columns of the matrix **A** of the AR parameters. When *λ*_2_=0 F-LASSO coincides with LASSO [26].

Finally, SG-LASSO is a convex combination of the LASSO and the group LASSO penalties [27]. This procedure imposes a structural constraint on the AR coefficients matrix, in addition to the basic sparsity one (*l*_1_ term), to model the assumption that the predictors can be aggregated in groups of a given size. In the penalty term of SG-LASSO, *g*is the number of groups, pre-determined, and 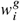 denotes the weight for the *i* − *th*group (in this study, as specified in [29], the weight of each group was set to 1). As [16] pointed out, in a multi-variate regression analysis, *M* groups of *p* elements for each column of **A** can be used. The *p* elements of each group, [*a*_*i j*_ (1), …, *a*_*i j*_ (*p*)] ∈ *ℜ*^*p*×1^ with (*i, j*) ∈ (1, …, *M*), can be represented by one AR parameter from the time lag 1 to time lag *p*. In other words, the second penalty term of the SG-LASSO selects a subset of groups by setting to zero all the coefficients in some groups (sparsity between groups), whereas the first term encourages sparsity within each group.

Since the different rows of the matrices **y**^*p*^ and **y** _*j*_ can be considered as independent from each other, it is possible to estimate the optimal value of lambda (*λ*_*opt*_) by dividing them into training and test sets. In the present work, 50% of the rows were used for the training set and 50% for the testing set. Training and test sets are standardized and the optimal values for *λ* are selected by using a Generalized Cross Validation (GCV) criterion [13, 18, 20, 30].

## COMPARING DIFFERENT PENALIZED LINEAR REGRESSION TECHNIQUES

This section reports the theoretical design and the practical implementation of a simulation of multiple interacting stochastic processes, which is used as a benchmark to illustrate the properties of the penalized regression techniques analyzed in this work. Using this simulation we show how the different techniques behave in different conditions of data paucity. Their performance is then compared by using different performance parameters.

### Simulation Study

Problem (3) was solved for the different penalty terms listed in Table I. We used the SLEP package (MatLab^®^) [29] with the multivariate linear regression solved with accelerated gradient methods [31, 32] by exploiting the parallel computing toolbox of MatLab. The code used in this paper, for the model identification through penalized regression techniques, is collected in the S-MVAR MATLAB toolbox available at https://github.com/YuriAntonacci/S-MVAR

The simulation study included the following steps:

1. Generation of simulated datasets, fitting predefined ground-truth networks under different values of the k-ratio (0.5, 0.8, 1, 1.5, 2, 3). The k-ratio is computed as the number of available data samples *N*_*s*_ divided by the number of parameters to be estimated *M p*.
2. Selection of the regularization parameters for each regression method by means of GCV criterion. The process was iterated 10 times for each *λ* randomly changing the training and testing sets.
3. Estimation of the AR parameters by using the six regression methods of interest, OLS, RR, LASSO, E-NET, F-LASSO and SG-LASSO.
4. Evaluation of the performances by comparing the estimated AR parameters with those imposed in the corresponding ground-truth network.

To increase the robustness of the statistical analysis, the entire procedure was repeated 50 times randomizing the data generation.

### Signal Generation

The simulated data-sets were generated according to different ground-truth networks by means of an MVAR model used as a generator filter [33]. The simulated multivariate time series (*M* = 10) were generated as realizations of a VAR(10) process fed by Gaussian noise with variance equal to 1. Furthermore, an autoregressive component extracted from a real EEG signal was on imposed in the model in order to reproduce its spectral properties [34]. The EEG signals used for this purpose were acquired from the *C*_*z*_ location of a healthy subject with sampling frequency equal to 200 Hz during the resting state. The simulated networks were randomly generated with a connection density of 15% (14 out of 90 possible connections). The matrix of AR parameters were automatically generated by the SEED-G software [34] by assigning randomly the lag in the range [1,10] and the coefficient value in the interval [-0.9, 0.9]. This value is imposed only at one lag among the possible *p* lags, randomly selected. The null connections are instead characterized by null coefficients at each lag. Under these constraints, 50 realizations of the VAR(10) process were generated for different values of the k-ratio parameter in the set {0.5, 0.8, 1, 1.5, 2, 3}, so that the length of the simulated time series was *N*_*s*_ = 50 when *K* = 0.5 and *N*_*s*_ = 300 when *K* = 3.

### Performances Evaluation

The performances of the investigated regression approaches were assessed in terms of accuracy in estimating the strength of the network link (i.e., values of MVAR coefficients) and in terms of ability to reconstruct the network structure. The error in the connection strength was evaluated separately for non-null and null links. Specifically, the values of the theoretical AR matrix **A** were compared with the estimated values stored in **Â** using the Mean Absolute Percentage Error (MAPE) [35] if the theoretical value is different from zero, and the Root Mean Squared Error (RMSE), otherwise:

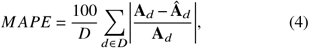

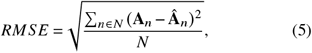

where *D* and *N* represent the set of non-null and null elements, respectively. The distributions of MAPE and RMSE across the 50 iterations were then presented separately for each method.

The ability in reconstructing the network structure was tested only for the subset of regression methods that yield a sparse matrix as output (i.e., LASSO, E-NET, F-LASSO and SG-LASSO). In particular, the problem of comparing the two AR parameter matrices, representative of the estimated and theoretical network structure, can be seen as a binary classification task. The existence (class 1) or absence (class 0) of each estimated connection is assessed and compared to the underlying ground-truth structure. Performances were assessed computing the false positive rate (FPR) that measures the fraction of null links for which an AR coefficient different from zero was detected, the false negative rate (FNR) that measure the fraction of non-null links for which an AR coefficient different from zero was detected, and the Area Under Curve (AUC) that summarized the information provided by FNR and FPR [36, 37]. These performance parameters were evaluated for each individual network and their distribution across the 50 simulated connectivity structures was then presented separately for each regression method.

The last considered performance parameter is the computational time (in seconds), required for the computation of the AR matrix **Â**. In particular, for penalized regressions, the estimation process is divided into two different steps: 1) selection of the regularization parameters, taking *T*_*sel*_ seconds; 2) computation of **Â** taking *T*_*comp*_ seconds. The process for the selection of the regularization parameters was performed by means of the GCV criterion within a range of 350 values for *»*_1_ and 100 values for *»*_2_ for each method and for each value of k-ratio (where applicable, see Table I). To speed-up the entire process, the parallel computing toolbox implemented in MatLab^®^ 2016a was used.

### Statistical Analysis

In the present work, a repeated measures two-way ANOVA test was performed for each performance parameter (MAPE, RMSE, FNR, FPR and AUC) in order to evaluate the effect of the factors: k-ratio with the levels {0.5, 0.8, 1, 1.5, 2, 3} (factor K); type of regression method with the levels OLS, RR, LASSO, E-NET, F-LASSO, SG-LASSO (factor TYPE). FNR, FPR and AUC were not considered in the ANOVA for the methods OLS and RR because they do not produce sparse AR matrices. The Greenhouse-Geisser correction for the violation of the spherical hypothesis was used in all the analyses. Tukey’s post-hoc test was used for testing the differences between sub-levels of ANOVA factors. Bonferroni-Holm correction was applied for multiple ANOVAs computed on different performance parameters.

### Results of the Simulation Study

The results of the two-way repeated measures ANOVAs computed separately for all the performance parameters are expressed in terms of F-values considering K and TYPE as within factors and reported in Table II (^* *^ is associated with *p* < 10^−5^). The two-way ANOVA performed on MAPE and RMSE reveals a strong statistical influence of the main factors K, TYPE, and their interaction *K* ×*TY PE* on the two performance parameters. Fig. 1 reports the distribution of the parameters MAPE and RMSE as a function of the interaction factor *K* × *TY PE*. The comparison of the six procedures shows remarkable differences between penalized regression techniques and the classical OLS. Specifically, MAPE decreases as the number of data samples available for the estimation procedure increases (Fig. 1.a). In particular, when K=3, there are no statistically significant differences between all methods, with a MAPE of approximately ∼ 20%. On the contrary, when *K* ≤ 1, OLS shows significantly higher values of MAPE (with a sharp increase between 100% and 280%) compared to penalized regression methods where MAPE values remain below 80%. Tukey’s post-hoc test reveals statistically significant differences between regression methods only for *K* = 0.5. In particular, MAPE computed using LASSO, was significantly lower than RR and F-LASSO.

**TABLE II.** F-values of the two-way repeated measures ANOVA.

| Factor | MAPE | RMSE | FNR | FPR | AUC |
| --- | --- | --- | --- | --- | --- |
| K | 367.2** | 324** | 140.6** | 23.7** | 143.7** |
| TYPE | 149.8** | 1863** | 85.3** | 277.5** | 129.1** |
| $K \times TYPE$ | 95.6** | 321** | 13.5** | 10.1** | 11.5** |

**FIG. 1.**
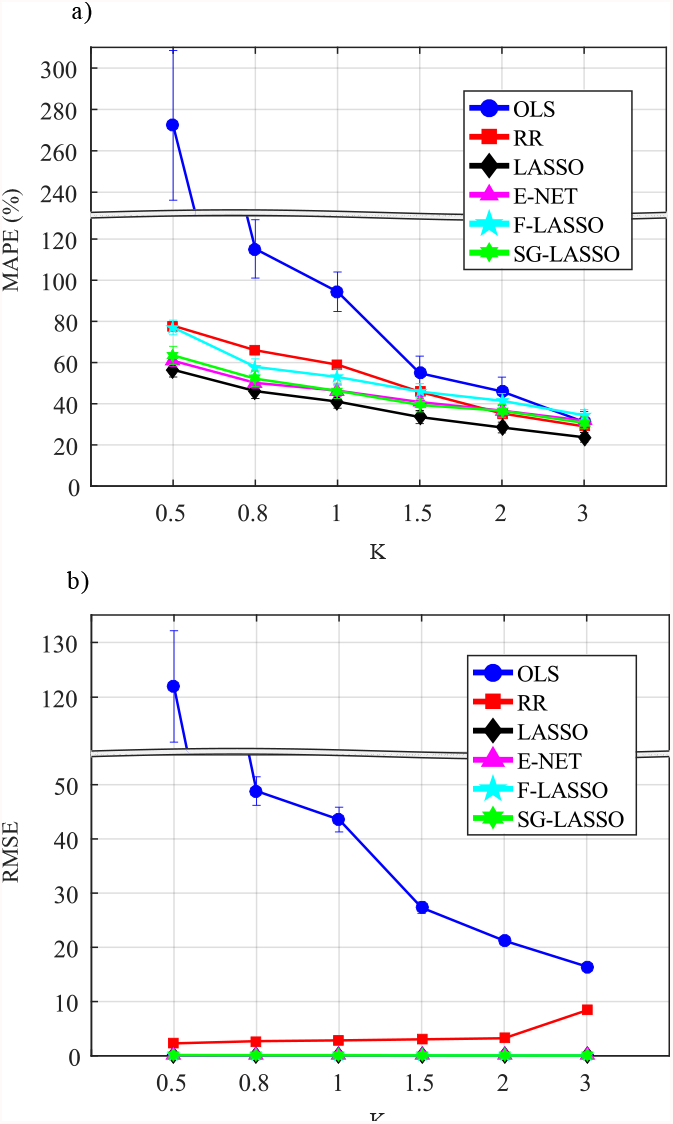
Performance parameters computed for non-null (MAPE, a) and null (RMSE, b) connections as a function of the factors K (k-ratio) and TYPE (regression methods). Both parameters are reported as mean value and 95 % confidence interval computed across 50 realizations for each experimental condition (6 regression methods and 6 values of K.

The analysis of RMSE (Fig. 1.b) reveals overlapping trends that are very close to zero, obtained for the penalized regressions LASSO, F-LASSO, SG-LASSO, and E-NET. Only RR exhibits a higher mean value of RMSE when compared with the other penalized regressions. However, it performs better than OLS. Similarly to the results obtained for the MAPE parameter, OLS shows a sharp increase in RMSE as the data samples (factor K) decrease. In this case as well, there is a noticeable discontinuity in the RMSE trend, between K=1.5 and K=1, as also highlighted in the analysis of MAPE.

The two-way ANOVA performed on FNR, FPR and AUC reveals a strong statistical effect of the main factors K and TYPE and of their interaction K × TYPE (Table II). Fig. 2 reports the distributions of the parameters FNR, FPR, and AUC as a function of the factor K for each regression approach performing a variable selection procedure (i.e., LASSO, E-NET, F-LASSO, SG-LASSO). All regression methods exhibit a decreasing number of false negatives as the number of available data samples increases (Fig. 2.a). However, under the most challenging condition of *K* = 0.5, F-LASSO, and SG-LASSO (azure line and green line, respectively) demonstrate the highest values of FNR, classifying roughly half of the theoretically non-zero links as null. LASSO and E-NET demonstrate better performance compared to all the other methods. No statistically significant differences are noticed between LASSO and E-NET, regardless of factor K, as highlighted by Tukey’s post-hoc tests. Also, LASSO and E-NET show false negative values ranging between zero (*K* = 3) and 30% (*K* = 0.5).

**FIG. 2.**
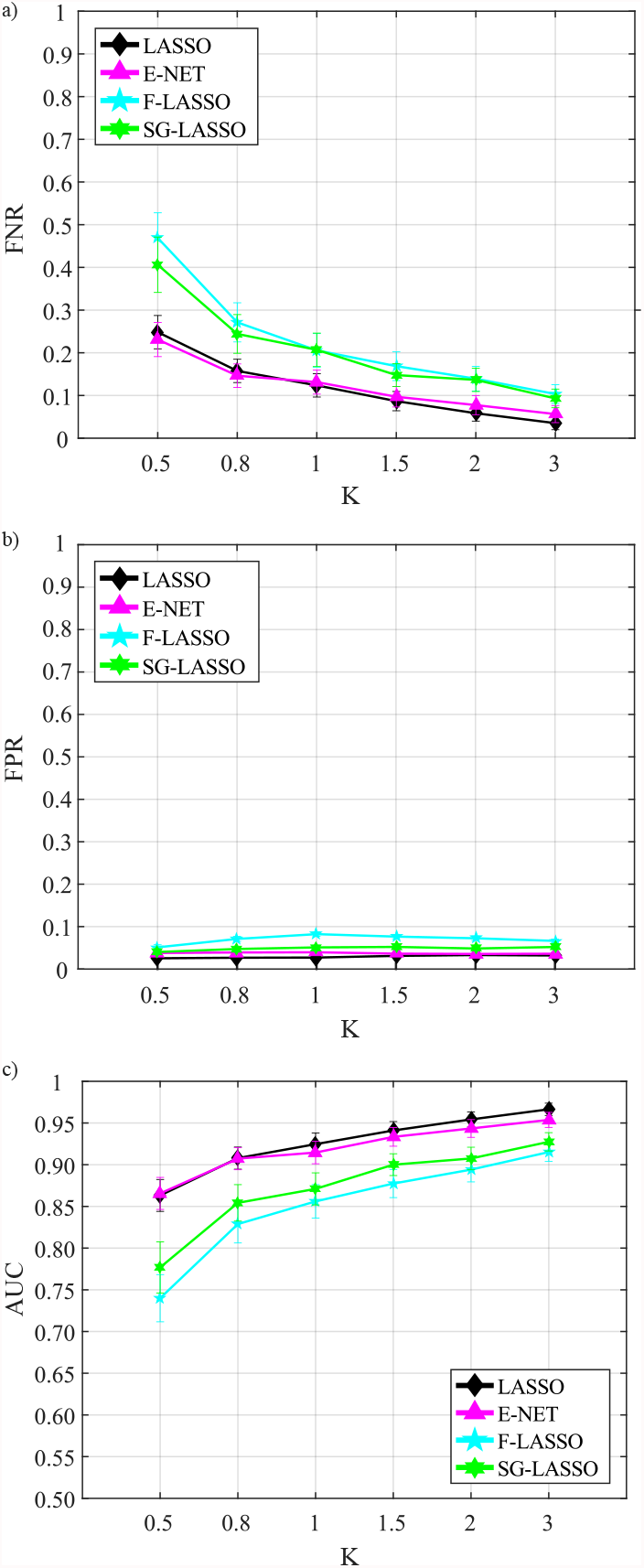
Distribution of FNR (a), FPR (b) and AUC (c) parameters considering the interaction *K* × *TY PE*, expressed as mean value and 95% confidence interval of the parameter computed across 50 realizations for the four regression methods, performing variable selection, and for different values of K.

The analysis of false positives (Fig. 2.b) reveals that the number of links incorrectly classified as non-null remains stable and almost negligible for all methods. The only exception is represented by F-LASSO which on average exhibits higher values of FPR compared to the other methods.

The overall network reconstruction performance, summarized by the AUC parameter (Fig. 2.c), demonstrates that LASSO and E-NET yield the most effective results. Specifically, when compared with F-LASSO and SG-LASSO, both LASSO and E-NET exhibit a higher rate of correctly detected links in all analyzed conditions. SG-LASSO performs better than F-LASSO, which shows the worst performance. For *K* = 3, LASSO and E-NET achieve an average AUC value close to 1, indicating a nearly perfect reconstruction of the network structure. Tukey’s post-hoc test indicates that there are no statistically significant differences between LASSO and E-NET. However, both LASSO and E-NET perform significantly better (higher AUC) compared to SG-LASSO and F-LASSO, irrespective of factor K. Additionally, AUC computed for SG-LASSO is significantly higher than F-LASSO. These regression methods demonstrate the ability to reconstruct the network structure with a very high accuracy of approximately 0.75 even in the worst-case scenario where the number of data samples is half the number of coefficients to be estimated.

Table III reports the computational time (in seconds) required for the selection of the optimal value of lambda/s (*T*_*sel*_) and the following computation of the MVAR parameters (*T*_*comp*_). The two values are computed for each regression method and for each value of K. In the case of OLS, it was not possible to evaluate the *T*_*sel*_ parameter since it is not included in the set of penalized regression methods. Times were recorded on a PC with an IntelCore i7-6700 processor, clock speed 3.40 GHz, 8 Gb RAM DDR4 (1.33 MHz), Intel (R) HD Graphics 530, 1024 Mb dedicated VRAM. Both, *T*_*sel*_ and *T*_*comp*_ increase with K. For all the values of K analyzed, the methods based on one single regularization parameter (i.e., LASSO and RR) need a shorter time for the selection step if compared with those based on two regularization parameters (i.e., F-LASSO, E-NET, and SG-LASSO). Furthermore, among the methods based on *l*_1_-norm, LASSO shows the lowest computational time required for the estimation process. The most time-consuming method is the SG-LASSO.

**TABLE III.** Computational time required for each regression method.

| Regression Method | $T_{sel}$ | | | | | | $T_{comp}$ | | | | | |
| --- | --- | --- | --- | --- | --- | --- | --- | --- | --- | --- | --- | --- |
|  | K=0.5 | K=0.8 | K=1 | K=1.5 | K=2 | K=3 | K=0.5 | K=0.8 | K=1 | K=1.5 | K=2 | K=3 |
| OLS | - | - | - | - | - | - | 0.03 | 0.02 | 0.03 | 0.05 | 0.05 | 0.06 |
| RR | 7.7 | 9.4 | 10.3 | 13.1 | 15.3 | 20.4 | 0.01 | 0.02 | 0.02 | 0.03 | 0.04 | 0.06 |
| LASSO | 6.7 | 10.8 | 12.9 | 19.2 | 23.8 | 35.7 | 0.02 | 0.03 | 0.03 | 0.04 | 0.05 | 0.06 |
| E-NET | 315 | 495 | 572 | 872 | 927 | 1400 | 0.03 | 0.03 | 0.04 | 0.04 | 0.05 | 0.09 |
| F-LASSO | 588 | 860 | 882 | 1200 | 1300 | 1700 | 0.05 | 0.06 | 0.06 | 0.07 | 0.08 | 0.09 |
| SG-LASSO | 870 | 1200 | 1500 | 1800 | 2000 | 2300 | 0.03 | 0.04 | 0.05 | 0.07 | 0.08 | 0.19 |

To summarize, all performance parameters are directly influenced by the amount of data samples available for the estimation process, exhibiting a direct proportionality. This means that the accuracy of the estimation process improves as the value of K increases. Even in cases where data is scarce, all the analyzed penalized regression methods demonstrate the ability to achieve good performance in both estimating the values of AR coefficients and reconstructing the network structure. Among the *l*_1_-norm-based regression methods, LASSO and EL-NET show the best performances, with no statistically significant differences between them. Notably, LASSO also exhibits a significantly shorter computational time when compared to EL-NET.

## TESTING OF LINEAR REGRESSION TECHNIQUES ON REAL EEG DATA

The study of human brain activity during motor imagery (MI) plays an important role in clinical neurology and neuroscience and for this reason the neuronal representation of MI and motor execution has been studied intensively for years using brain imaging techniques such as functional magnetic resonance [38], EEG [39] and positron emission tomography [40]. It is well established that adults present 8-12 Hz (i.e. *μ*) and 16-26 Hz (i.e. *β*) rhythms in the EEG recorded from the scalp over the primary sensorimotor cortex. These rhythms show amplitude fluctuations that are synchronized to movement imagery [41]. These fluctuations are more prominent in the contralateral primary motor cortex and contralateral somatosensory regions as demonstrated in [42]. Given their clearly defined occurrence, the information derived from these oscillations and their localization on the scalp has also been used for developing brain computer interfaces [43]. Since MI involves a dynamic interplay between separate brain regions, it is reasonable to assume that brain connectivity can provide useful information for understating this phenomenon. For instance, in the context of brain computer interfaces, the feasibility of using brain connectivity as an additional feature for the discrimination between different mental tasks has been explored [21, 44]. However, the authors pointed out how the accuracy during the entire process of connectivity estimation drops dramatically when few data samples are available and it would be necessary to use penalized regressions to improve the classification accuracy. In this section, by using the EEG dataset previously used in [21], we demonstrate that it is possible: (i) to estimate brain connectivity using penalized regression techniques with a limited amount of data samples, and (ii) to discriminate between two different experimental conditions by relying on features extracted from a brain connectivity analysis.

### Data description and pre-processing

The EEG dataset made available by the authors of [21] consists of 45 EEG channels located over the scalp according to the international 10-20 system. The signals were recorded at a sampling rate of 300 Hz with three synchronized g.USBamp amplifiers (g.tec, Guger Technologies OEG, Graz, Austria) and pre-processed following the pipeline described in [21]. To test the regression methods on real EEG signals, a healthy subject (male, right handed), with no prior experience in brain computer interface (BCI) control based on motor imagery, was selected to record one session, consisting of 90 trials of right hand motor imagery (HAND) and 90 trials of foot motor imagery (FOOT). Further details about the experimental paradigm and the pre-processing process are available in [21].

### Single-trial Connectivity Estimation

To reproduce the condition of the simulation study, 11 out of the 45 available channels were selected (*C*_5_, *C*_3_, *C*_1_, *C*_2_, *C*_4_, *C*_6_, *CP*_3_, *CP*_4_, *C*_*z*_, *CP*_*z*_, *FP*_*z*_) and, as suggested in [21], 100 samples between the third and the fourth second were selected to obtain a proper time window in which the task was correctly performed. Each of the eleven time series obtained from each trial and condition (HAND -FOOT) were interpreted as a realization of a VAR process whose matrix of parameters Â was estimated with the six different regression methods (i.e. OLS, RR, LASSO, EL-NET, F-LASSO, SG-LASSO). The order of the model, denoted as *p*, was estimated for every experimental condition and trial. This estimation was done using the Final Prediction Error (FPE) criterion [1]. Afterwards, the average value of the model order across trials and conditions was calculated, resulting in a value of 10. All the analyses were performed by identifying MVAR models of dimension *M p*, where M=11 and with the model order fixed to *p* = 10, which together with a time series of 100 points guarantees a connectivity estimation with a value of K close to 1 for each regression method.

To retrieve information about the connectivity between EEG signals in the frequency domain, the estimated AR parameters were then Fourier transformed [10].

### Classification Task

To verify (i) if penalized regressions can be used for estimating connectivity between EEG time series and, (ii) if the features derived from a connectivity analysis can provide useful information even in data paucity conditions, a classification task was performed.

The classification HAND vs. FOOT was repeated by using as features the frequency version of estimated AR parameters (real part of the Fourier transform of the AR parameters) in two different frequency bands, typically related to the MI tasks [43]: *α* (8-12 Hz) and *β* (13-30 Hz). By considering *f*_*i*_ as the frequency interval under investigation and *M* = 11, it was possible to extract *M*^2^ *f*_*i*_ = 605 features in *α* band (*f*_*i*_ ∈ [8, 12]) and 2178 in *β* band (*f*_*i*_ ∈ [13, 30]).

In order to ensure a ratio of at least 5 between the number of training cases and the number of classifier parameters to be estimated, i.e., to avoid overfitting during the training phase, a sub-selection of the features was performed [45]. In particular, we selected the top 25 highest -ranked ones according to the associated t-value (independent samples t-test, HAND vs FOOT) for each feature in each frequency band [46]. Note that all the diagonal elements of the AR matrices were removed to maintain only information related to causal effects.

The data were divided into 70% for the training process (126 cases out of 180), 15% for validation (27 cases out of 180) and 15% for testing (27 cases out of 180). Each of the three data sets contained the same number of observations for each class (HAND-FOOT) to avoid class imbalance problem [47]. By using a holdout approach at each iteration, one different feed-forward neural network (FFNN) was trained, validated, and tested [45]. The structure chosen for the FFNN is one of the most widely used for classification purposes [48, 49] that includes one hidden layer with one neuron (sigmoid activation function) and an output layer with a softmax activation function [50]. The initial weights of the network were randomly generated and the training was performed with a gradient descent algorithm (with learning rate set to 10^−3^), with cross-entropy used as the cost function [51]. The training process was stopped by means of the early stopping criterion [52]. The process was repeated 100 times for each considered frequency band. As a performance parameter, we computed the classification accuracy (ACC) on the test set [53].

For this study, a two-way repeated measures ANOVA test on the classification accuracy (ACC) was performed in order to evaluate the performance of classification accuracy depending on the regression method (factor TYPE: OLS, RR, E-NET, LASSO, F-LASSO, SG-LASSO) and the frequency band (factor BAND: *α,β*). Moreover, to assess statistical differences between pairs of distributions independent samples t-tests were performed.

### Results on real EEG data

Fig. 3 reports the distributions of the classification accuracy evaluated on the test set according to the interaction *BAND* × *TYPE* for each regression method and for *α* and *β* frequency band.

**FIG. 3.**
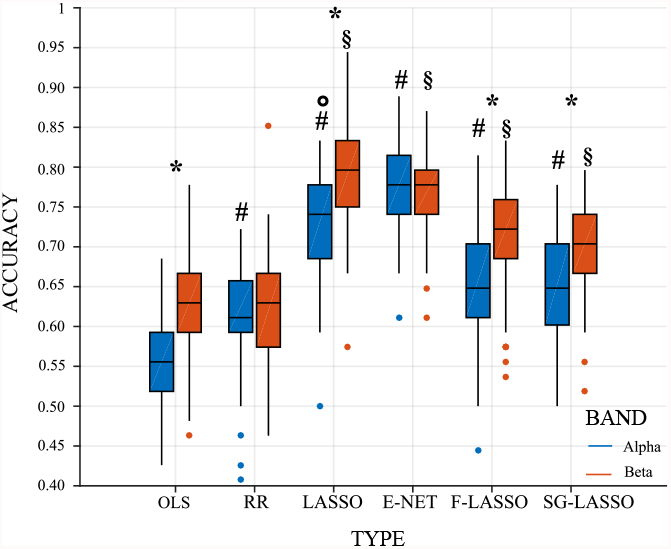
Classification accuracy evaluated on the testing set for each regression method obtained for each frequency band (*α* and *β*). Box plots report the distribution of the classification accuracy across the 100 iterations by considering the interaction *TYPE* × *BAND* (F=18.57, *p* 10^−5^). On each box, the central line indicated the median, the edges of the box indicate 25^*th*^ and 75^*th*^ percentiles, and outliers are marked with a circle. Statistically significant differences between pairs of distributions are marked with * (*α* vs *β*), with # (· vs OLS for *α* band), with § (· vs OLS for *β* band), and with º (LASSO vs E-NET in *α* band)

For all the regression methods, except RR and E-NET, ACC is significantly greater in *β* band (blue boxes) with respect to *α* band (orange boxes). The analysis of *α* band reveals a statistically significant increment of the ACC if penalized regressions are used when compared with OLS (distributions marked with #); this is not the case for the *β* band in which RR is not significantly different from OLS. All the other regression methods (i.e. LASSO, E-NET, F-LASSO, and SG-LASSO) show a statistically significant increment of ACC with respect to that obtained with OLS (distribution marked with §). LASSO and E-NET show the highest values of ACC (∼ 0.8) but only using LASSO it is possible to highlight a significant difference between *α* and *β* frequency bands. On the other hand, OLS shows the worst performance in discriminating between HAND and FOOT with an average value of ACC ∼ 0.55 in *α* band and ~ 0.65 for *β* band.

Table IV reports the time required for estimating the AR coefficients matrix Â for each regression method. Also in this case the process of cross-validation described in Section III.C was used for the selection of different optimal values of lambda/s with the same PC used for the simulation experiment. As a first observation, the computational times required are comparable to those obtained in the simulation study (see Table III) performed on *M* = 10 EEG-like data for the case *K* = 1. Table IV shows that OLS, which does not require the selection of a regularization parameter, is the fastest method. Among penalized regression methods, RR and LASSO show a difference of a few seconds for the selection of *λ*_*opt*_ (∼ 7*s*, ∼ 8*s* respectively). E-NET, F-LASSO, and SG-LASSO represent the slowest regression methods with several minutes needed for the selection of *λ*_1_ and *λ*_2_ (∼ 12 *min* for E-NET and ∼ 32 *min* SG-LASSO).

**TABLE IV.** Computational time required for each regression method (*K* ∼ 1)

| Regression Method | $T_{sel}$ | $T_{comp}$ |
| --- | --- | --- |
| OLS | - | 0.04 |
| RR | 6.9 | 0.05 |
| LASSO | 8.1 | 0.11 |
| E-NET | 719 | 0.15 |
| F-LASSO | 997 | 0.16 |
| SG-LASSO | 1978 | 0.18 |

## DISCUSSION

The present work compares the performances of different regression techniques for the study of connectivity estimates in the framework of linear signal processing. The behavior of the different regression techniques was validated both in a simulation setting and in a real-world scenario relevant to the study of brain connectivity from EEG signals.

### Simulation experiment

The simulation study was designed to assess the performance of the classic OLS when compared with five different penalized regression techniques (RR, LASSO, E-NET, F-LASSO, and SG-LASSO) used in the identification procedure of the MVAR model. Performance parameters were evaluated in terms of the strength of the network links (values of the MVAR coefficients) as well as network structure (false positives and false negatives). Lastly, the computational time needed to solve all the linear problems was analyzed.

The accuracy in estimating the value of the MVAR coefficients was investigated across different k-ratio values by means of MAPE and RMSE used as performance parameters (Fig. 1). As expected, both performance parameters for all the regression methods show a tendency to increase as the k-ratio decreases. This tendency is more evident for OLS, as already documented testing different VAR identification approaches (e.g., the Levinson recursion for the solution of Yule-Walker equations) in the context of signal processing [12]. For K=1 the uniqueness of the solution is not guaranteed for OLS as the matrix ([y*p*^*T*^y^p^), necessary for the solution of the problem (2), approaches singularity. As a consequence, OLS exhibits a strong bias, as reported in previous studies [13, 24, 54]. On the other hand, we document very low values of MAPE and RMSE for penalized regressions, even when K≤ 1, confirming their high tolerance to collinearity between regressors caused by the drastic reduction of data samples available [13, 55]. The trends obtained for MAPE highlight the best performances of LASSO and E-NET without any statistically significant differences between them.

The ability in the reconstruction of the network structure was investigated in terms of false negatives, false positives and AUC. The latter appears to be the best-suited indicator in terms of its capability to synthesize the similarity of two networks also in the condition of class imbalance, a typical condition in sparse networks, as those simulated in this work [36]. The regression methods tested here are those based on *l*_1_-norm which in turn induces sparsity within the matrix of estimated MVAR coefficients [14]. Focusing on Fig. 2, all the penalized regressions are characterized by a low percentage of false positives (FPR ∼ 0.05). Additionally, both false negatives and AUC values increase as K decreases. In particular, the maximum value of AUC (∼ 0.95) is obtained by LASSO and E-NET regardless of the value of K. Even in the worst condition (*K* = 0.5), these two methods reach high values of AUC (above 0.85). On the other hand, F-LASSO showed the worst performance for all the values of K, with AUC∼ 0.75 when K is equal to 0.5. The results obtained in this study align with those reported in a previous work that utilized Ridge and LASSO regression to investigate brain connectivity estimated from fMRI data [20]. Specifically, the authors demonstrated how penalized regression techniques can effectively identify causal relationships, even when the number of nodes in the network exceeds the available data samples. The obtained AUC values ranged from 0.6 to 0.9. Moreover, these findings are consistent with our previous work, which compared only OLS and LASSO in the context of information dynamics analysis [13].

The observed differences between LASSO and E-NET when compared to other penalized regressions can potentially be explained through a methodological consideration. In their work, Zou et al. (2003) [25] highlighted that LASSO and EL-NET may not be the ideal approaches when dealing with structurally grouped regressors (referred to as **y**^*p*^ in this study). Specifically, only SG-LASSO introduces a structural constraint to account for grouped variables, for instance when the predictors are represented by genes. In the study of connectivity, where prior information about potential connections between different brain areas is lacking, it is reasonable to assume that LASSO and EL-NET may exhibit superior performance in retrieving the theoretical network structure compared to SG-LASSO and F-LASSO.

Finally, the computational time required from each method to provide an estimate of the coefficient matrix Â was computed for each regression method. Two different computational times were reported, one for the GCV procedure used for the selection of the penalization terms (depending on whether the methods present one or two parameters for regularization) and the other one for the MVAR parameters computation (Table II). The total time required for the entire estimation procedure increases as the value of K increases. As expected, regression methods that incorporate two regularization parameters (E-NET, F-LASSO, and SG-LASSO) require more time because all combinations of the two parameters need to be tested through GCV. The less time-consuming regression methods are LASSO (among those based on *l*_1_ norm) and the E-NET regressions (among those based on a linear combination of *l*_1_ and *l*_2_-norm). To limit the computational time, a smaller number of values could be tested for the different regularization parameters (i.e. *λ*_1_, *λ*_2_). However, this could have an impact on the performance of the selected regression method since the optimal solution may not be in the range selected by the operator. It is therefore recommended to use at least a range of 100 values for each regularization parameter.

### Application of linear regression techniques on real EEG data

A well-established body of literature demonstrates an intrinsic balance between excitatory and inhibitory coupling among brain regions and between hemispheres occurring in the execution of a motor imagery task [56]. Based on this evidence, we selected the EEG time series recorded from a total of 11 channels including the left hemisphere (contralateral to the hand motor imagination), the right hemisphere (ipsilateral), and the midline. As a consequence, our EEG results are discussed in terms of primary motor cortex activity of the contralateral and ipsilateral hemispheres [57].

The simulation study showed that penalized regressions are reliable tools for estimating brain connectivity in conditions of data paucity where the classical OLS fails. Previous studies highlighted the possibility of discriminating different experimental conditions by using features related to brain connectivity, such as the estimated AR parameters and their frequency version [58, 59]. However, it was pointed out that, with the current methodology based on the OLS estimator, it is impossible to reach an appropriate accuracy and it is necessary to move towards penalized regression techniques [21]. The combined use of features derived from brain connectivity with neural network classifiers showed an increase in classification accuracy [60] with most of the results found in *α* and *β* frequency bands [39, 59]. Interestingly, the results reported here show comparable classification accuracy even for different tasks and for a different number of data samples available for the estimation procedure. In fact, as reported in Fig. 3, LASSO and E-NET attain the highest values of ACC in the *α* and *β* frequency bands (∼ 0.8). Furthermore, as proof of the unsuitability of OLS in a condition of strong data paucity (*K* ∼ 1), Fig. 3 showed the lowest values of ACC in both frequency bands.

## CONCLUSIONS AND IMPLICATIONS

The aim of this work was to evaluate the usefulness of penalized regression methods for brain connectivity estimation. The simulation study showed how LASSO and E-NET can estimate, with high accuracy, not only the value of AR coefficients but also the related connectivity structure even in conditions of data paucity in which OLS fails (e.g. when collinearity between regressors arises for the lack of data points).

The results consider the possibility of discriminating between two tasks through a classifier trained with features extracted from a brain connectivity analysis. LASSO and E-NET showed the best performances in terms of accuracy of classification. Notably, these results were obtained for *K* = 1, a condition in which OLS fails whereas E-NET and LASSO showed comparable performances when simulated EEG signals were used. These findings suggest that, when OLS cannot be used, LASSO might represent the most suitable alternative, even when a limited amount of computational power is available.

If confirmed on a larger sample, the overall results pave the way for using sparse identification procedures for connectivity estimation in all those conditions in which few data points are available, such as in the estimation of brain networks at the single trial level as well as during real-time applications. Since all the linear connectivity estimators are based on the identification of an MVAR model, it is reasonable to assume that penalized regressions could be used and integrated for the computation of all the connectivity estimators in both time and frequency domains as already partially demonstrated [61, 62]. Future developments will aim at testing penalized regressions for information dynamics analysis or for the study of high-order dependencies between time series extracted from complex systems dynamics using the VAR modeling approach [63, 64].

## Notes

### Competing Interest Statement

The authors have declared no competing interest.

